# Deciphering Intercellular Communication in the Cerebellar External Granule Layer: Insights into Non-Classical Connections in Neural Development

**DOI:** 10.1101/2025.08.11.669634

**Authors:** Malalaniaina Rakotobe, Shiyu Liu, Garima Virmani, Gabriel Kaddour, Nathaly Dongo Mendoza, Frédéric Doussau, Jean Livet, Laurence Cathala, Chiara Zurzolo

## Abstract

Intercellular communication is essential for brain development. While classical modes—paracrine, juxtacrine, and synaptic signaling—are well characterized, emerging evidence suggests that membranous bridges, such as tunneling nanotubes (TNTs), forming de novo between cells and mainly described *in vitro*, may also contribute. Yet their presence and function in vivo remain unclear, partly due to the difficulty of distinguishing them from other intercellular connections (ICs). Building on connectomic observations in fixed tissue reporting ICs in the external granule layer (EGL) of the developing cerebellum, we examined their nature in postnatal day 7 (P7) mice. Using immunofluorescence, sparse genetic labelling, and live imaging, we distinguished division-independent ICs from cytokinetic bridges (CBs) and intercellular bridges (IBs). CBs were detected in the EGL, whereas IBs were not observed in this region. In addition to CBs, we identified membranous protrusions which appeared to link clonally and non-clonally related cells. The presence of these ICs reinforces previous connectomic evidence in fixed tissue and raises the possibility that they may participate in intercellular communication during cerebellar development. These findings warrant further investigation into a potentially underexplored mode of communication in the developing brain.

## Introduction

Intercellular communication plays a crucial role in brain development, ensuring the precise regulation and coordination of processes including neurogenesis, cell migration, differentiation and the establishment of functional neuronal networks. This communication occurs through adhesion molecules, the secretion of extracellular vesicles and soluble molecules (Bahram Sangani et al., 2021; Sahay et al., 2020), as well as through synaptic interactions via electrical and chemical synapses (Pereda, 2014). Disruptions in these pathways can lead to neurodevelopmental disorders, highlighting their significance in brain maturation.

Beyond classical mechanisms, recent studies have identified additional structures involved in intercellular communication, including cytonemes, intercellular bridges (IBs), and tunneling nanotubes (TNT) (Korenkova et al., 2020). Cytonemes are thin, F-actin-based, close-ended protrusions (1–700 μm in length) that mediate signal transduction between cells via membrane-tethered receptors and ligands, without establishing cytoplasmic continuity (Hall et al., 2024). IBs (0.4–350 μm in length) arise from incomplete cytokinesis and allow cytoplasmic exchange between connected cells (Haglund et al., 2011; Rujano et al., 2022).

Unlike cytokinetic bridges (CBs), which are abscised at the end of cytokinesis, IBs persist longer, facilitating sustained communication between connected cells. TNTs, which can extend up to 200 μm, are F-actin based, open-ended membranous structures, formed *de novo* between two distinct cells. They facilitate the transfer of cargo such as organelles, proteins, and nucleic acid. The open-ended nature of TNTs can be assessed using high-resolution electron microscopy (EM) or live imaging which demonstrates uninterrupted transfer of large cargo such as organelles (Sáenz-de-Santa-María et al., 2023). However, these techniques are difficult to apply to tissue samples. Identifying TNTs in *vivo* is particularly challenging due to the lack of specific molecular markers, the fragility of these structures, and their sensitivity to fixatives like paraformaldehyde, which can compromise structural preservation. Consequently, such structures are typically referred to as “TNT-like” in tissue until their ability to transfer cellular cargos between connected cells is confirmed (Palese et al., 2025). Due to these hurdles, TNTs have been extensively studied *in vitro*, but their *in vivo* characterization remains limited, with only recent evidence confirming functional TNTs in the zebrafish embryo (Korenkova et al., 2024).

In the developing cerebellum, the external granule layer (EGL) constitutes a dynamic region where granule cells (GCs), the most numerous neurons of the brain, are generated. The EGL is subdivided into an outer proliferative zone (oEGL) and an inner differentiating zone (iEGL), as originally defined by Altman & Bayer (Altman & Bayer, 1997). In rodents, granule cell progenitors (GCPs) proliferate in the oEGL for approximately three weeks before differentiating into postmitotic GCs, which migrate through the iEGL and molecular layer (Curran & O’Connor) to reach the inner granular layer (IGL) (Consalez et al., 2020). Intercellular communication within the EGL is essential for coordinating interactions not only between GCs but also with other neuronal populations (Kim et al., 2023). Combining serial-section scanning electron microscopy (ssSEM) with computational connectomics in fixed tissue, we recently identified open-ended intercellular connections (ICs) linking immature GCs in the mouse EGL. By analyzing cellular parameters such as shape, Golgi apparatus number, and cilium presence, our data suggested that these structures could represent either stable IBs or TNTs. However, their precise nature and function could not be precisely determined since the ssSEM approach did not allow the use of specific markers to distinguish IBs from other types of cellular connections (Cordero Cervantes et al., 2023).

In this study, we examined the presence and characteristics of ICs in the EGL of postnatal day 7 (P7) mice. Using immunofluorescence, live imaging, and sparse genetic labelling, we differentiated cytokinetic bridges (CBs) and intercellular bridges (IBs) from other connections that appeared to span fully between cells. While CBs were identified in the iEGL, our data indicate that they do not account for all previously described ICs (Cordero Cervantes et al., 2023), suggesting that other types of connections—such as TNT-like structures—may also occur during cerebellar development.

## Materials and methods

### Animals

All mice used were generated and maintained in the animal facility of the Institut Pasteur, Paris. The care and handling of the animals prior or during the experimental procedures followed European Union rules and were approved by the French Animal Protection Committee and the local Ethics Committees. Animals were maintained on 12h light/dark cycles with food and water provided *ad libitum*. C57BL/6 mice at P7 were used for the quantification of CBs and mitotic cells in the iEGL. Rj:SWISS mice were used for the electroporation of the *pCAG-Dendra2-E2A-EGFP-Cep55* plasmid.

### Transgenic mouse lines

We used a *CAG-Cytbow* reporter line bearing a *Cytbow* MAGIC Marker transgene (Loulier et al., 2014) that stochastically expresses mTurquoise2, eYFP or tdTomato under the broad *CAG* promoter in a Cre-dependent manner (this line will be described elsewhere). To generate experimental animals, we crossed homozygous *CAG-Cytbow^+/+^* males with *Nestin-CreER^T2+/-^*females (Lagace et al., 2007) to obtain *Nestin-CreER^T2+/-^; CAG-Cytbow^+/-^* offspring. To increase the probability of combinatorial fluorescent protein expression of in a Cre-dependent manner, these *Nestin-CreER^T2+/-^; CAG-Cytbow^+/-^* females were backcrossed with *CAG-Cytbow^+/+^* males to generate *Nestin-CreER^T2+/-^; CAG-Cytbow^+/+^*experimental animals.

### Tamoxifen injection

To sparsely label neural cells, *Nestin-CreER^T2+/-^; CAG-Cytbow^+/+^* newborn mice received subcutaneous injection of 25 µL of tamoxifen (234-118-0, Merck) diluted in corn oil once per day at P0, P1, P2 and P3 at a concentration of 5 mg/mL.

### Plasmid

Coding sequence of *pCAG-Dendra2-E2A-EGFP-Cep55* construct was cloned from *pDEST-Dendra2-E2A-EGFP-Cep55* construct (used in Korenkova et al., 2024) into the *pCAG-GFP* plasmid from Addgene (Plasmid #11150).

### Postnatal electroporation

Vetergesic, at a concentration of 0.05 mg/kg, was injected 30 minutes prior to surgery for analgesia. Anesthesia was induced with 4% isoflurane for 10 minutes and then maintained at 2% throughout the procedure. Before making the incision, laocaine at a concentration of 2.5 mg/kg was injected at the incision site to alleviate pain. To prevent desiccation, the eyes were protected with Ocry-gel. Following the craniotomy, the *pCAG-Dendra2-E2A-EGFP-Cep55* plasmid was injected at the surface of the cerebellum using a glass micropipette. The plasmid was delivered at 4–5 sites for a total injection volume of 1 µl at a concentration of 5.9 µg/µl. Triple electrodes coated with conductive gel were used for electroporation. The electrode connected to the anode was placed on a micromanipulator and lowered into contact with the craniotomy under optical control, while the two cathode electrodes were positioned on either side of the animal’s head. Electroporation (BTX electroporator) was performed with the following parameters: 5 pulses of 50 ms at 1 Hz, at 80V. Immediately after electroporation, the mouse was placed on a heating pad and stimulated to restart respiration in case of cardiorespiratory arrest. The incision was sealed with Vetbond surgical glue, and a subcutaneous injection of 20-30 μl of 0.9% NaCl was administered gently to rehydrate the mouse.

### Live imaging of acute cerebellar slices

This protocol was adapted from a Star Protocol by Qu. X et al. (Qu et al., 2021). Briefly, pups were decapitated and the brain dissected out in oxygenated, ice-cold complete HBSS (cHBSS) buffer. The brain was then transferred to the slicing chamber of a vibratome (Leica Biosystem VT 1000S), which contained cHBSS to keep the brain submerged, while oxygenation is maintained in the chamber. Cerebellar slices (250 μm thick) were prepared using the vibratome at a speed of 0.08–0.1 mm/s and collected in a “nest beaker” containing complete cHBSS medium, bubbled at 33°C. Cerebellar slices expressing fluorescent reporters were selected and transferred to a µ-Slide 8 Well (ibidi, Cat. No: 80807) with 1 mL of warm, fresh, sterile medium. The ibidi well were placed in a temperature-controlled chamber attached to the microscope, where both temperature (37°C) and CO□ flow (5%) were regulated and maintained to promote cell viability and ensure optical stability during live imaging.

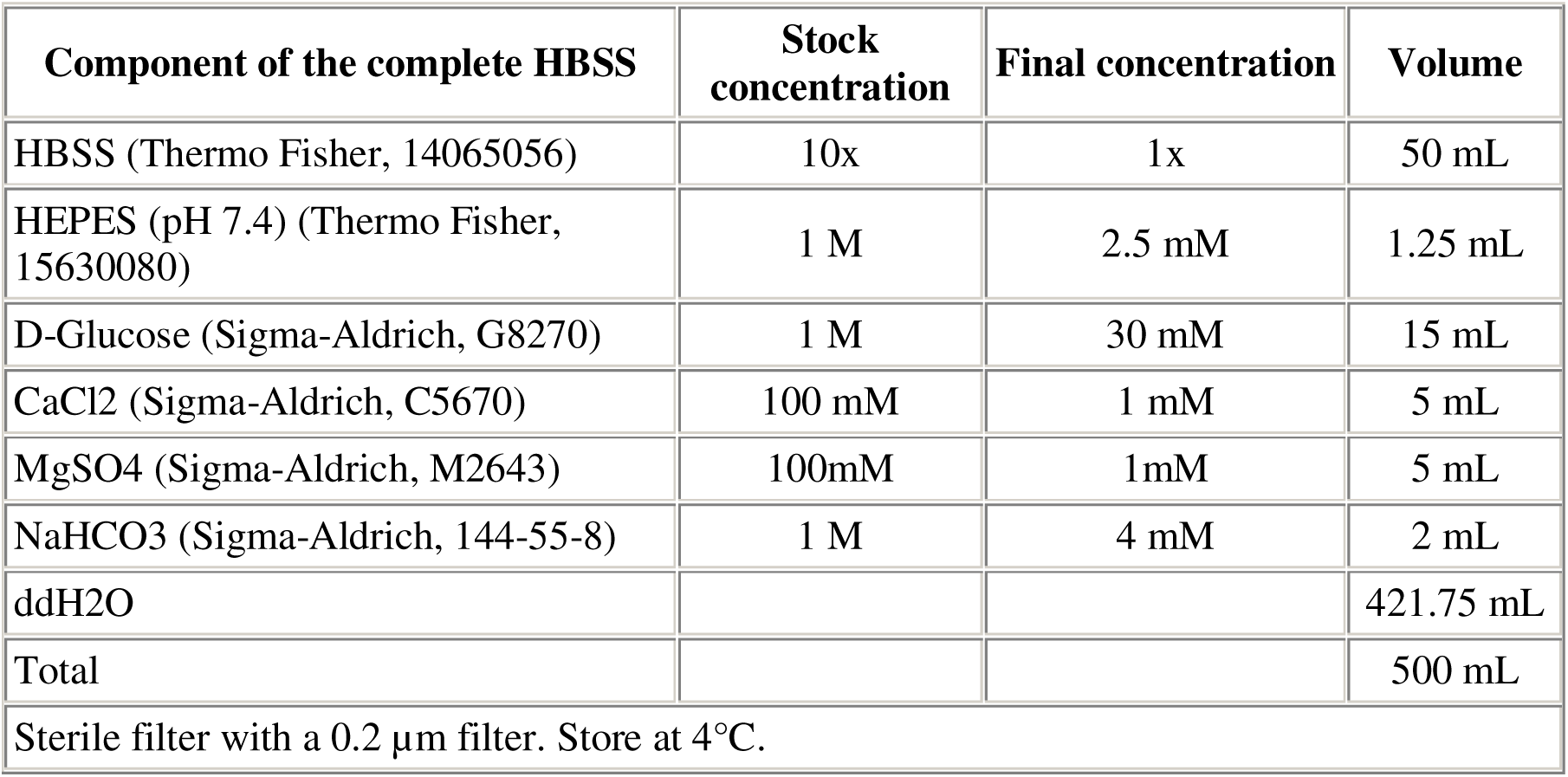

| Component of the complete HBSS | Stock concentration | Final concentration | Volume |
| --- | --- | --- | --- |
| HBSS (Thermo Fisher, 14065056) | 10x | 1x | 50 mL |
| HEPES (pH 7.4) (Thermo Fisher, 15630080) | 1 M | 2.5 mM | 1.25 mL |
| D-Glucose (Sigma-Aldrich, G8270) | 1 M | 30 mM | 15 mL |
| CaCl<sub>2</sub> (Sigma-Aldrich, C5670) | 100 mM | 1 mM | 5 mL |
| MgSO<sub>4</sub> (Sigma-Aldrich, M2643) | 100mM | 1mM | 5 mL |
| NaHCO<sub>3</sub> (Sigma-Aldrich, 144-55-8) | 1 M | 4 mM | 2 mL |
| ddH<sub>2</sub>O |  |  | 421.75 mL |
| Total |  |  | 500 mL |
| Sterile filter with a 0.2 $\mu\text{m}$ filter. Store at 4°C. | | | |

### Immunohistochemistry

Brains were dissected out and fixed overnight in 4% paraformaldehyde (PFA) at 4°C, then rinsed in PBS and cryoprotected in 30% sucrose in PBS at 4°C overnight before embedding and freezing in OCT (Cryomatrix™, Thermo) at −80°C. The cerebellum was sliced in sagittal sections of 40 µm using a Leica CM 3050S cryostat. Sections were then mounted on SuperFrost+ slides (Fisher Scientific). Slices underwent an antigen unmasking step: they were incubated in unmasking solution (citrate buffer, pH 6) for 15 minutes in the 2100 Antigen Retriever. They were then cooled in PBS for 10 minutes before being permeabilized for 10 minutes in PBS containing 0.5% Triton. Next, slices were incubated for 1 hour in blocking solution (PBS with 5% fetal calf serum (FCS) and 2% bovine serum albumin (BSA)). To quantify CBs and mitotic cells, two rabbit-derived primary antibodies were used sequentially (the rabbit anti-Aurora B kinase antibody and the rabbit anti-Ki67 antibody). For those samples a two-step IHC was run. First, slices were immuno-labelled with the rabbit anti-Aurora B primary antibody before running a second IHC with the other antibodies including the rabbit anti-Ki67 antibody. The first Rabbit antibody was included in the blocking solution at room temperature for 2 hours. After PBS washes, the slices were incubated overnight at 4°C with a Fab fragment secondary antibody in blocking solution. Then the second step of the IHC procedure was the same for all samples. Primary antibodies were incubated overnight at 4°C in blocking solution. The next day, secondary antibodies were incubated for 2 hours at room temperature in blocking solution. Finally, sections were incubated with DAPI (1 µg/ml) in PBS for 30 minutes and mounted using Fluoromount-Gold. Primary antibodies used were: rabbit anti-Aurora B kinase (Abcam, ab2254, 1:500), rabbit anti-Ki67 (Abcam, ab15580, 1:200), goat anti-TAG1 (rndsystems, AF4439, 1:200), mouse anti-citron kinase (BD Biosciences, 611376, 1:200), goat anti-anillin (Abcam, ab5910, 1:50), rabbit anti-anillin (Bethyl laborories, A301-406A, 1:500), chicken anti-GFP (Aves Lab, GFP-1010, 1:1000), goat anti-tdTomato (Sicgen, AB8181, 1:200), rat anti-PH3 (Abcam, ab10543, 1:200).

Secondary antibodies used (Jackson ImmunoResearch, 1:500) were: Fab Fragment anti-rabbit Alexa Fluor™647, anti-rabbit Alexa Fluor™594, anti-rabbit Alexa Fluor™647, anti-goat Alexa Fluor™488, anti-mouse Alexa Fluor™488, anti-goat Alexa Fluor™647, anti-chicken 488, anti-goat Alexa Fluor™594, anti-rat Alexa Fluor™488.

### Microscopy and image analysis

#### Imaging

Images of cerebellar sections were acquired on a Zeiss LSM900 confocal microscope, except for a subset of samples electroporated with the *pCAG-Dendra2-E2A-EGFP-Cep55* plasmid which were imaged on a Nikon Ti2E Spinning disk microscope. In addition, a Leica SP8 confocal microscope was used to image the 4 fluorescent proteins (eBFP2, mTurquoise2, eYFP, tdTomato) expressed by *Nestin-CreERT2+/-; Cytbow+/+* mice.

The Zeiss LSM900 was equipped with a 63x/1.4 NA oil-immersion objective (**Fig. 1, 2, 4, S4E**) or a 10x/0.45 NA DRY objective (**Fig. 3B, S3 and S4D**), as indicated on the figure legends. Images were acquired at 16-bit depth, with a pixel size of 0.1 µm and a z-step size of 0.35 µm. The following fluorochromes and detection settings were used: DAPI (excited at 405 nm, emission detected at 415-480 nm), Alexa Fluor 488 (488 nm, 500-550 nm), Alexa Fluor 594 (561 nm, 580-620 nm), and Alexa Fluor 647 (633 nm, 640-700 nm).

**Fig. 1.**
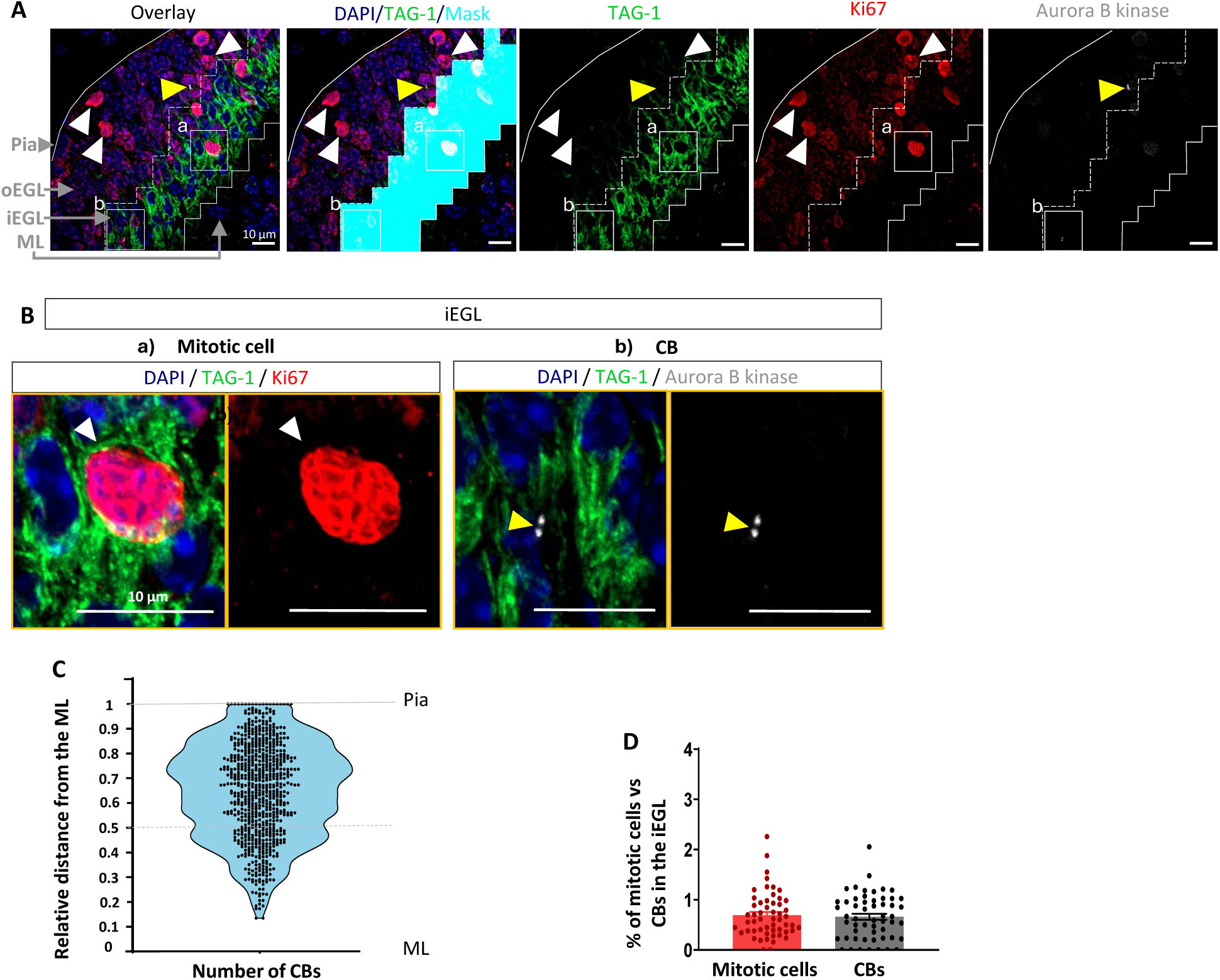
Assessment of the presence of CBs in the iEGL. (A) Sagittal sections of P7 WT cerebellum, taken from the medial vermis (lobule VIII), labelled with anti-TAG-1 (green), anti-Ki67 (red), anti-Aurora B kinase (grey) and DAPI staining (blue). TAG-1 and Ki67 expression were used to distinguish the iEGL from the oEGL using a CNN-based classifier that automatically generated a mask for the iEGL; the resulting boundary is indicated by the dotted line. Solid lines delineate the EGL, with the pia located above and the ML below. Yellow arrowhead points at a CB and white arrowheads at mitotic cells in the oEGL. (B) Higher magnification views of the regions shown in (A) highlighting (a) a mitotic cell (white arrowheads) and (b) a CB (yellow arrowhead) within the iEGL. (C) Distribution of CBs across the EGL, represented as the relative distance from the ML to the pia (0 = ML, 1= pia). The oEGL-iEGL transition corresponds approximately to the mid-point of this axis. (D) Comparison of the percentage of mitotic cells (0.69%) and CBs (0.66%) in the iEGL. The data are presented as mean ± SEM; p-value= 0.7 analyzed using a Student T-test. (C-D) N = 4 mice, 55 images were analyzed in total. All images are maximum-intensity projections of six z-slices acquired at 63x magnification.

**Fig. 2.**
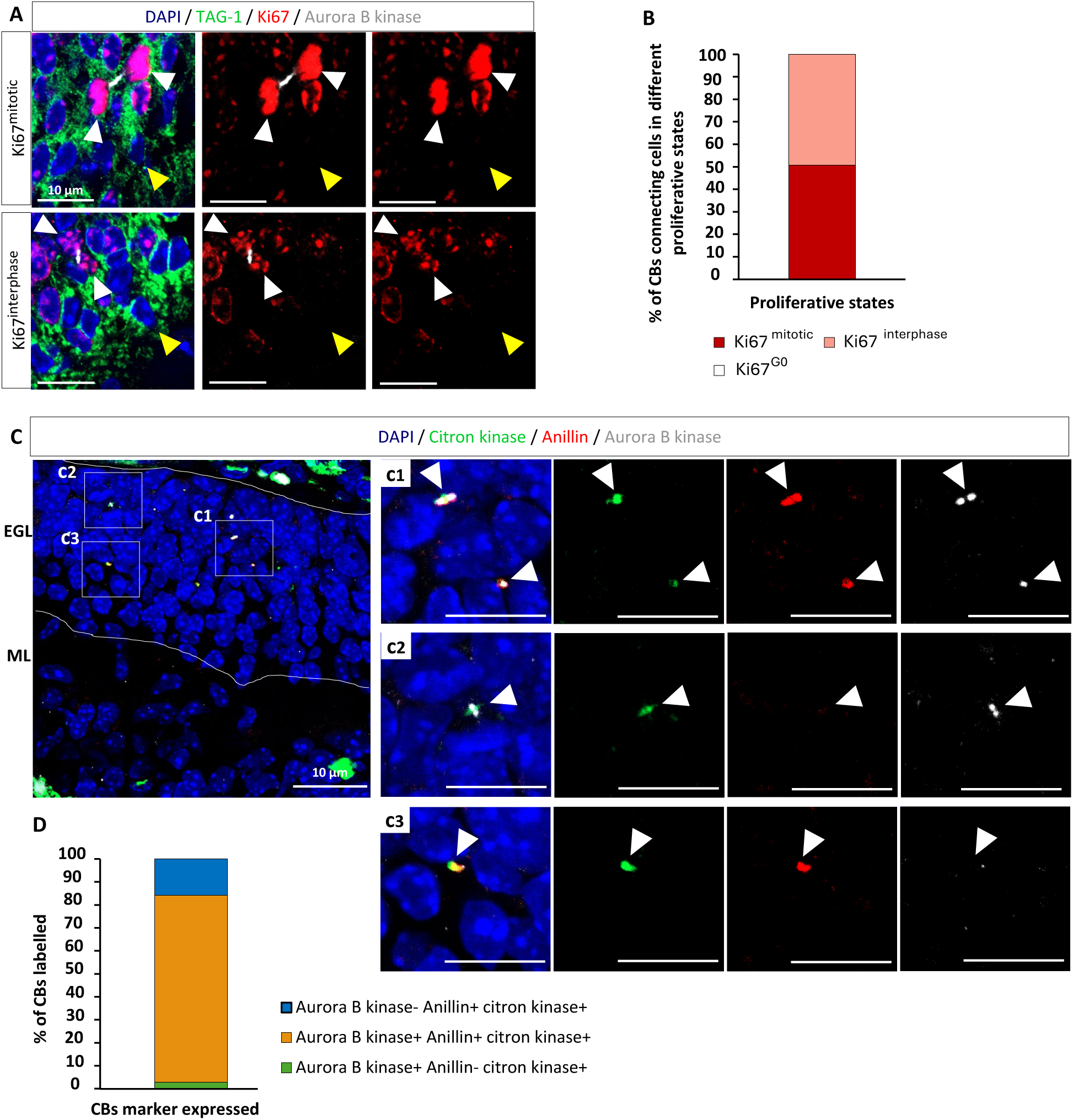
Assessment of the nature of CBs in the iEGL. (A) Sagittal sections of WT P7 cerebellum, taken from the medial vermis (lobule VIII), labelled with anti-TAG-1 (green), anti-Ki67 (red), anti-Aurora B kinase (grey) and DAPI staining (blue). The expression of TAG-1 was used to identify the iEGL using a CNN-based method that creates a mask on the iEGL area. The white arrowheads show Ki67+ cells that are connected by a CB. The yellow arrowheads show Ki67-cells, which were never observed to be connected by a CB. (B) Distribution of CBs connecting cells in the different proliferative states in the iEGL. The proliferative state is determined based on the pattern and expression level of Ki67. “Ki67^mitotic^” (51%) represents cells that are undergoing mitosis, “Ki67^interphase^” (49%) represents cells in earlier cell cycle phases or those that have recently exited proliferation, “Ki67^G0^” (0%) represents cells in late G0. Per-image percentages were averaged across all images from each mouse to obtain a mean value per animal. N = 4 mice; 55 images were analyzed in total. (C) Labelling with anti-citron kinase (green), anti-anillin (red), anti-Aurora B kinase (grey) and DAPI staining (blue). (c1-c3) Higher magnification of the areas indicated in (C), showing CBs labelled with both anillin and Aurora B kinase (c1), only Aurora B kinase (c2) or only anillin (c3). All CBs expressed citron kinase. (D) Proportion of CBs in the categories shown in c1-c3: 81.33% were Aurora B kinase^+^/Anillin^+^/citron kinase^+^, 2.78% were Aurora B kinase^+^/Anillin^-^/citron kinase^+^, and 15.89% labelled Aurora B kinase^-^/Anillin^+^/citron kinase^+.^ N = 4 mice, 66 images were analyzed in total. Pannel (A) corresponds to a z-slice, (C) and (c1-c3) are maximum projections of 6 z-slices (z-step=0.35 µm). All images were acquired at 63x magnification.

**Fig. 3.**
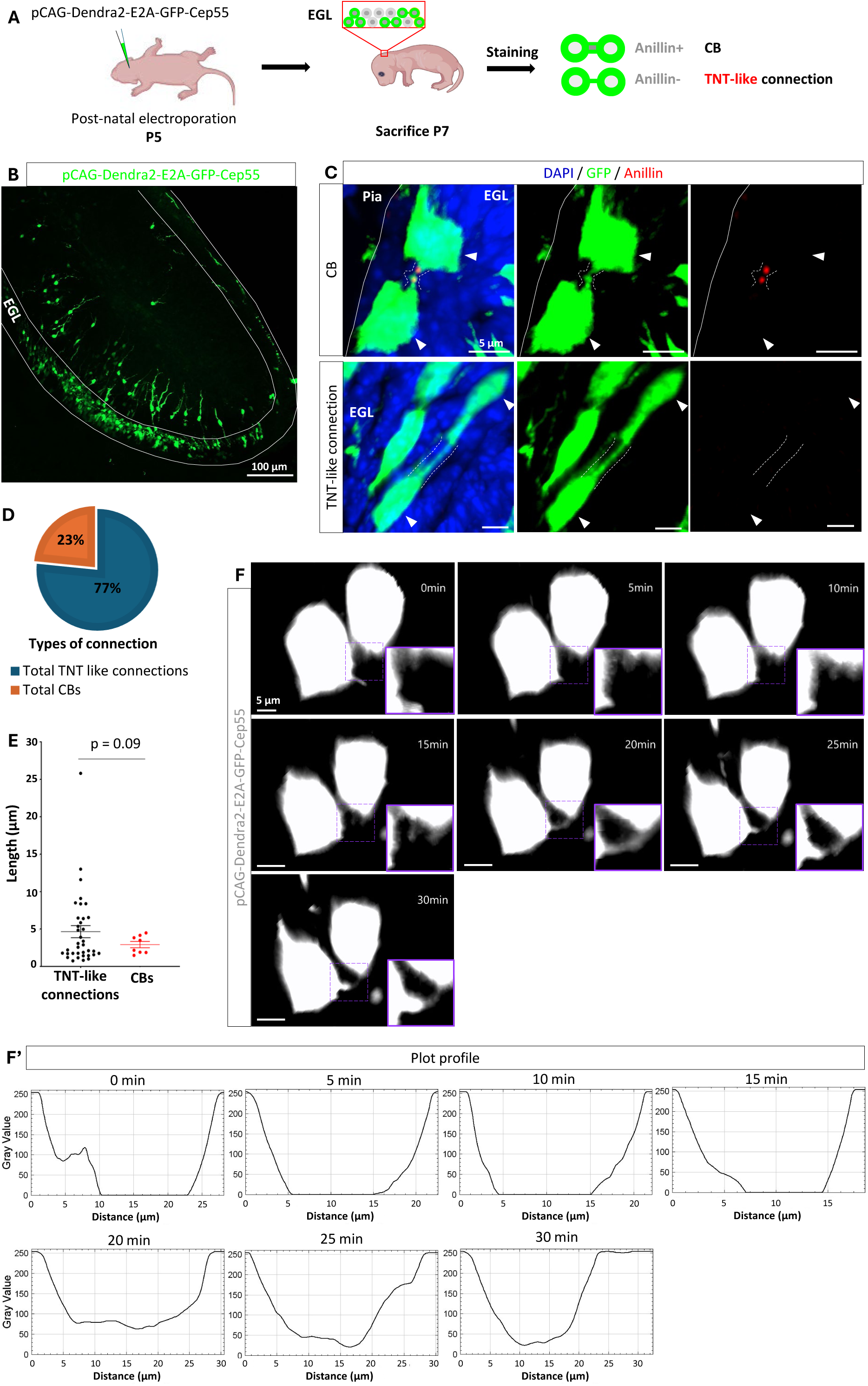
Identification of potential GC connections compatible with TNT-like bridges. (A) Schematic representation of the strategy used to sparsely label cells and visualize ICs: The pCAG-Dendra2-E2A-GFP-Cep55 plasmid is electroporated in the EGL of P5 pups. The pups are sacrificed at P7 and cerebellar slices were stained with anti-anillin to distinguish CBs from putative connections non-derived from cells division (ie. putative TNT-like connections). (B) Representative image showing the expression of the plasmid in the EGL. (C) Sagittal cerebellar slices labeled with anti-GFP (green), anti-anillin (red), and DAPI, showing a CB (anillin^+^ connection) and a TNT-like connection (anillin-connection). The white arrowheads point at the connected cells. Dotted lines delineate the connections. (D) Graph representing the proportion of CBs and TNT-like connections over the total number of ICs identified. (E) Length of the connections identified in (D) showing an average TNT-like connections and IB’s length of 5.38 µm and 2.92 µm, respectively. (D-E) N = 3 mice, 36 TNT-like connections and 11 CBs identified, 83 images were analyzed in total. Although cells were labelled, no connections were found in the EGL of one mouse, likely due to the rarity of these connections. In (E) data are presented as mean ± SEM; p-value= 0.09 analyzed using a Welch T-test. (F) Example of TNT-like connection forming between two cells, via the filopodia driven mechanism in a sagittal cerebellar slice. A zoomed-in view of the area where the connection forms (indicated by the dotted purple square) is shown in the bottom right corner of each time point. The two protrusions establish connection approximately 20 min after the start of the recording. (F’) Plot profile of the connection shown in (F) at the different time points, showing a continuous connection (grey value > 0) between 20-30 min. Panel (B) shows a z-maximal projection at x10 magnification. (C,F) images correspond to 3D views at 60x magnification. Solid lines delineate the EGL.

**Fig. 4.**
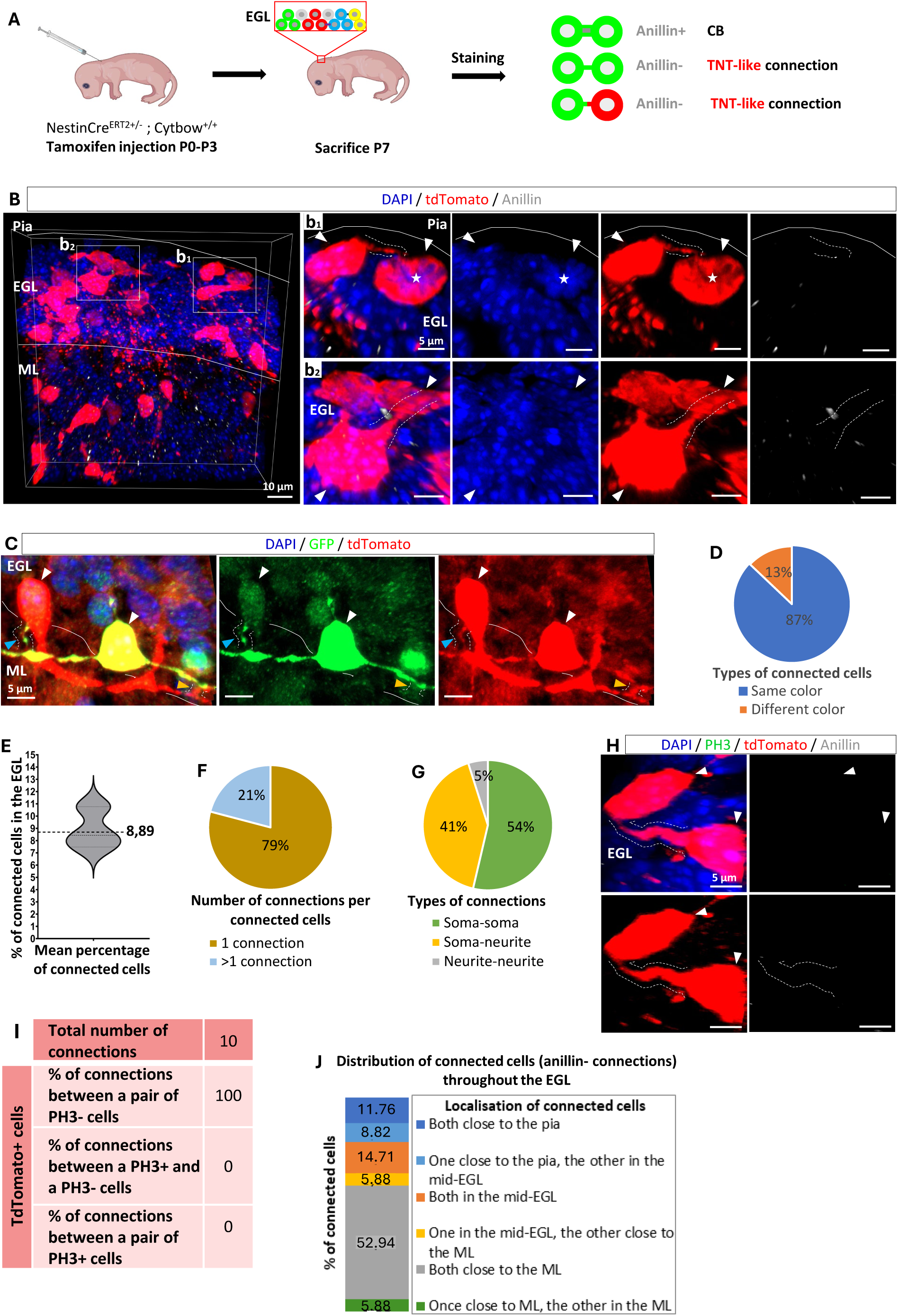
Assessment of the features of potential connections in the EGL. (A) Schematic representation of the strategy used to sparsely label cells and visualize ICs: *NestinCre^ERT2+/-^; Cytbow^+/+^* mice were injected with tamoxifen between P0 and P3 to induce the mosaic expression of mTurquoise2, eYFP, tdTomato or a combination of two of these reporters in neurons. Mice were sacrificed at P7 and were stained with DAPI, anti-anillin (grey) to distinguish CBs from putative TNT-like connections, anti-GFP (green) and anti-tdTomato (red) to amplify reporter signals and differentiate clonally vs. non-clonally related cells. (B) Representative image of TNT-like connections (anillin-) between tdTomato+ cells in the EGL. (b_1-_ b_2_) Represent a higher magnification of the areas indicated in (B) showing TNT-like connections close to the pia (b_1_) and in the middle of the EGL (b_2_). The asterisk marks a dividing cell, as identified by the DAPI pattern. (C) Representative image showing a TNT-like connection between a tdTomato^+^ cell and a tdTomato^+^GFP^+^ cell near the ML border. The blue arrowhead points at a connection formed between the soma and neurite. The orange arrowhead points at a connection between the neurites. (D) Graph presenting the percentage of TNT-like connection between same-color vs. different-color cells. (E) Graph representing the relative percentage of connected cells over the total labelled population. The dotted line indicates the mean. SEM is ± 0.061. (F) Graph presenting the percentage of cells forming one (53 out of 67 GCs) vs. multiple (>1) TNT-like connections (14 out of 67 GCs). (G) Graph presenting the percentage of TNT-like connections categorized by location: soma-soma (22/41 connections), neurite-neurite (17/41 connections), soma-neurite (2/41 connections). (H) Labelling of EGL sagittal sections with anti-PH3 (in green), anti-Tdtomato (in red), anti-anillin (in grey) and DAPI staining (in blue). (J) Spatial distribution of connected (I) Quantification representing the percentage of TNT-like connections (anillin^-^) between non-dividing cells (PH3^-^), dividing cells (PH3^+^), one dividing (PH3^+^) and one non-dividing cell (PH3^-^) cells across the EGL. (b_1_-b_2,_ C, F, I) The white arrowheads indicate connected cells. The dashed lines delineate TNT-like connections. (D-E, G-H, K) N = 3 mice, 200 images. (I-J) N = 3 mice, 86 images. All images correspond to 3D views and were taken at 63x magnification. Solid lines delineate the EGL. Dotted lines delineate the connections.

The Nikon Ti2E inverted microscope was equipped with a Yokogawa CSU-W1 spinning disk unit and a Photometrics Prime 95B back-illuminated sCMOS camera. A 60x/1.4 NA oil-immersion objective was used (**Fig. 3C, 3F**). Images were collected in 12-bit mode, with a pixel size of 0.11 µm and a z-step of 0.3 µm. Standard bandpass filters optimized for each fluorophore were used including 447/60 for DAPI and 525/50 for AF488.

The Leica SP8 confocal microscope was used with either a 20x/0.75 NA water immersion objective (**Fig. S4B**) or a 63x/1.4 oil-immersion oil objective (**Fig. S4C**), as indicated in the figure legends. Images were collected in an 8-bit mode for low-magnification tile scans (**Fig. S4B**) and in 16-bit mode for high resolution acquisition (**Fig. S4C**). For the 20x objective, the pixel size was 1.14 µm with a z-step of 3.8 µm. For the 63x objective, images were single tiles with a pixel size of 0.09 µm and a z-step of 0.35 µm. A 405 nm laser and white laser (470-670) were used to excite the following fluorophores: eBFP2 (excited at 405 nm, emission detected at 415-465 nm), mTurquoise2 (excited at 470 nm, emission detected at 480-530 nm), mEYP (excited at 514 nm, emission detected at 524-574 nm), tdTomato (excited at 554 nm, emission detected at 564-604 nm).

All image quantifications were performed on unprocessed datasets. Brightness and contrast adjustments applied to figures were linear and applied across entire images, solely to enhance visualization of thin TNT-like structures.

#### Quantification of mitotic cells

Connected nuclei were assigned to one of three Ki67 pattern classes based on the nuclear distribution of the Ki67 signal: (i) Ki67^mitotic^, showing a dense, compact/perichromosomal pattern characteristic of mitotic cells; (ii) Ki67^interphase^, displaying a punctate or dotted nuclear pattern typical of interphase; and (iii) Ki67^G0^, lacking detectable nuclear Ki67 signal. These categories follow published descriptions of Ki67 dynamics across the cell cycle (Sobecki et al., 2016; Sun & Kaufman, 2018). Scoring was performed manually on z-stacks using DAPI as a nuclear reference. For each image, only mitotic cells connected by a CB were quantified.

#### Normalization of mitotic cells and CB quantification in the iEGL

Quantification of CBs and mitotic cells in the iEGL were normalized to the total number of nuclei within the EGL to allow comparison with our previous study (Cordero Cervantes et al., 2023). Nuclei were segmented using a multi-step computational pipeline combining deep learning and classical segmentation. First, a U-Net convolutional neural network was trained to segment nuclei in 3D confocal stacks (**Fig. S2A**). The resulting binary masks were then used to remove background signals.

Next, nuclear detection was refined in 2D using Cellpose (Stringer et al., 2021) which proved more reliable than the previously used watershed algorithm (**Table 1**). To estimate the number of nuclei in 3D, Cellpose-based 2D counts were corrected using a scaling factor (F□) (F□ = 0.092) determined by comparing automatic 2D counts to manual 3D annotations on representative stacks. The correction factor Fi for each region was calculated as below and applied uniformly to all images:

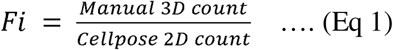

**Table 1.** Estimation of a 2D-to-3D correction factor for Cellpose-based nucleus counting. Correction factors were derived by comparing Cellpose 2D counts to manual annotations of the same slice (Manual 2D count) and to the full Z-stack (Manual 3D Count). Relative Error quantifies the discrepancy between Cellpose 2D and manual 2D counts. The average Correction Factor (F□)=0,092 was then used to extrapolate total 3D counts from Cellpose 2D count across the full dataset.

| Sample | Region | Cellpose 2D<br>count | Manually count | | Relative<br>error (%) | Correction<br>Factor ($F_i$)<br>(according to<br>Eq. 1) |
| --- | --- | --- | --- | --- | --- | --- |
|  |  |  | 2D | 3D |  |  |
| Sample 1 | Region n°1 | 547 | 503 | 56 | 8.75% | 0.11 |
| Sample 2 | Region n°2 | 735 | 692 | 62 | 6.21% | 0.09 |
| Sample 3 | Region n°3 | 695 | 668 | 62 | 4.04% | 0.09 |
|  |  |  |  | Average | 6.33 % | 0.092 |

#### Nuclei segmentation

Nuclei segmentation with the U-Net CNN (VGG-16 encoder) involved four sequential steps: data preparation, learning, testing, and inference. The DAPI channel was used as input; slices were rescaled (0-255), equalized, and manually annotated in VAST (Berger et al., 2018) to generate ground-truth masks.

Training data were split into overlapping 1×320×320-pixels tiles (Z, X, Y), with a stride of 1×270×270 pixels. Pixel dimensions were calibrated to 0.35□µm/pixel in X and Y axes, and the Z-spacing between slices was 3.5□µm. All image tiles were rescaled before training and split randomly into training (80%) and test (20%) sets.

Augmentations included random rotations, flips, ±10% translations, and ±20% contrast adjustments. The U-Net model was trained for 40 epochs using the Adam optimizer (initial learning rate: 1×10□□, batch size 2) and selected based on the best intersection-over-union (IoU) score.

During inference, the trained model was applied to the full 3D volumes using a sliding-window approach with the same tile size and overlap used during training. For each tile, probability maps were thresholded at 0.5 to produce binary masks, stitched, and post-processed by hole filling and border-object removal to refine the final segmentation. The resulting masks were used to isolate nuclei and eliminate background signals.

### EGL and iEGL classification

The iEGL was delineated using a CNN-based classifier (was the same as described in our previous paper (Cordero Cervantes et al., 2023)) trained on three-channel inputs (DAPI, Ki67, TAG1; **Fig. S2B**). Input volumes were split into non-overlapping patches of 100×100 pixels (X, Y) (0.35 µm/pixel). The network comprised three convolutional blocks with increasing filter sizes (16, 32, 64), each followed by max-pooling, and a dense layer of 128 units with dropout (rate 0.2). Input images were resized to 100×100 pixels, and pixel values were normalized to the [0, 1] range. The training dataset consisted of randomly chosen and manually labeled iEGL and Background tiles. A total of 5488 random tiles were used for training and 1372 for validation. To enhance model generalization, data augmentation was applied during training, including random horizontal flips, ±10% rotations, and ±10% zoom.

For iEGL class after filtering: 2658 tiles (Training: 2126, Test: 532) For Background class after filtering: 8404 tiles (Training: 6723, Test: 1681) The final model achieved F1-scores of 0.99897541 for iEGL and 0.99833055 for background on the test data. During inference, patches with probability > 0.5 for iEGL and < 0.6 for background were classified as iEGL, and binary masks were reconstructed into full 2D maps. The resulting classified patches were reconstructed into binary masks delineating the iEGL. To generate the full EGL mask, the post-processed iEGL mask was expanded by padding, with the padding direction (left, right, up, down, right-up, or left-down) manually selected depending on the anatomical orientation of the EGL in the image (**Fig. S2C**).

The number of nuclei in the iEGL was determined by segmenting nuclei in 2D and counting those that overlapped with the iEGL mask. To estimate the corresponding 3D nuclear count, the 2D count was multiplied by the previously derived scaling factor F□=0.092 (**Fig. S2D**). The same approach was used to quantify the nuclei in the EGL (**Fig. S2D’**).

#### Image analysis and quantification

Analyses in **Fig. 1** and **Fig. 2** were performed on sections from the medial vermis (lobule VIII), consistent with our previous study (Cordero Cervantes et al., 2023). Analyses in **Fig. 3** and **Fig. 4** were conducted across the entire cerebellum, although labelled cells were predominantly located in lobule IX and X. 2D images were processed with Fiji (ImageJ). 3D renderings of acquired z-stacks were generated using Imaris® software (version 10, Oxford Instruments). TNT-like connections were identified based on morphological criteria: a membranous protrusion was counted as a TNT-like connection if it appeared continuous, did not express CBs or IBs markers, and connected two labelled cells. The total number of labelled cells was counted manually using the Spots function in Imaris. The length of every TNT-like connection was measured using the Measurement points tool.

All quantifications were performed on confocal images acquired from multiple cerebellar sections per mouse. Counts of labeled structures were normalized per image to the total number of DAPI□ nuclei within the relevant region (EGL or iEGL), defined using CNN- and Cellpose-based segmentation for CBs and mitotic cells, or by manual counting of the total number of labeled cells for TNT-like connections. For **Fig.1** and **Fig.2**, per-image percentages were averaged across all images from each mouse to obtain a mean value per animal. For **Fig. 4**, the percentage of TNT-like connections was calculated by summing all connections and labeled cells across images for each mouse, then averaging these values across all mice.

Percentages within categories (Ki67 classes, Aurora B□ and/or anillin□ bridges) were calculated per image and averaged per mouse as described above. N refers to the number of m ice analyzed, and data are reported as mean ± SEM across biological replicates.

Statistical analysis was performed using Microsoft Excel and GraphPad Prism. To compare relative numbers of CBs and mitotic cells Student’s T-test, as the variance was equal and the data were normally distributed. Illustrations of the quantifications are displayed as violin plot, dot plot, bar, and pie chart. Each experiment was performed in at least 3 biological replicates.

## Results

### Cytokinetic bridges (CBs) are present throughout the EGL but less frequent in the iEGL

To assess the presence of mitotic cells and CBs in the EGL in P7 mice, we labeled mitotic cells with an anti-Ki67 antibody and CBs with an anti-Aurora B kinase antibody (**Fig. 1A, S1A**). Expression of the Transient Axonal Glycoprotein TAG-1, together with Ki67, was used to distinguish the iEGL from the oEGL through a CNN-based approach that automatically generated a mask of the iEGL region. In the oEGL, mitotic cells were found in high density and displayed a distinct and robust Ki67 expression pattern (**Fig. 1A**) (Sobecki et al., 2016; Sun & Kaufman, 2018) consistent with previous reports identifying this layer as the primary proliferative zone where cell division is most frequent (Legué et al., 2016; Nakashima et al., 2015)—and consequently CB formation. The iEGL contrasted with a sparse distribution of mitotic cells (**Fig. 1A, B, white arrowheads)**, indicating that only a small number of GCs continue to divide within this layer. This was corroborated by anti-Aurora B kinase labelling, which presence is indicative of CB at the end of mitosis (**Fig. 1A, 1C** yellow arrowhead) with a progressive decrease in CB number toward the ML (**Fig. 1C0**) (N = 4 mice; 55 images, 615 CBs quantified across the EGL). As expected, their proportion in the iEGL was also lower compared to the oEGL (**Fig. 1C**). These findings align with previous studies characterizing the EGL as a dynamic proliferative zone (Cordero Cervantes et al., 2023; Hanzel et al., 2019). Moreover, our quantification of mitotic cells is consistent with our previous finding, despite different methodological approaches (Cordero Cervantes et al., 2023) (**Fig. S1A)**.

To determine whether the previously identified intercellular connections (ICs) (Cordero Cervantes et al., 2023) could be fully accounted for by CBs, we replicated our previous analysis by quantifying mitotic cells (prophase, prometaphase, metaphase, anaphase, and telophase) and Aurora B kinase–positive CBs in the iEGL using immunofluorescence, and compared these values with those previously obtained using an ssSEM-based approach (**Fig. 1D, S1A, S1B**). All quantifications were normalized to the total number of nuclei within the EGL, obtained from CNN- and Cellpose-based segmentation of the DAPI channel (see **Fig. S2** and Methods). Immunolabeling marked similar percentages of mitotic cells (0.69%) and CBs (0.66%) in the iEGL, consistent with ongoing mitotic activity in this layer and suggesting that the labeled structures likely represent transient post-mitotic intermediates. However, the frequency of CBs obtained with this approach was substantially lower than the overall frequency of ICs (3.76%) previously reported in our earlier study using ssSEM (Cordero Cervantes et al., 2023) (see **Fig. S1A** for a comparison), which suggests that additional types of connection, such as IBs or TNT-like connections, may also be present in the iEGL.

### The EGL exhibited no detectable intercellular bridges (IBs)

To assess whether IBs are present in the iEGL at P7, we performed quantitative immunofluorescence to evaluate the proliferative state of GCs connected by Aurora B kinase-positive bridges. Cell-cycle classification was based on Ki67 nuclear distribution patterns rather than absolute intensity (see Methods), enabling discrimination among 3 stages (**Fig. 2A**): mitosis (Ki67^mitotic^), interphase (Ki67^interphase^), and quiescence (Ki67^G0^) (Sobecki et al., 2016; Sun & Kaufman, 2018). Connected GCs were nearly equally distributed between the “Ki67^mitotic^” (51%) and “Ki67^interphase^” (49%) classes (**Fig. 2B).** In most cases, both connected cells exhibited comparable Ki67 patterns, consistent with their shared mitotic origin. Occasionally, asymmetric Ki67 labeling was observed within a pair, likely reflecting early post-mitotic divergence or initiation of cell-cycle exit in one daughter cell. However, Aurora B kinase-positive CBs were never observed between pairs of “Ki67^G0^” GCs (**Fig. 2A, yellow arrowhead, and 2B).** These findings indicate that CBs can persist transiently following mitotic exit, but do not remain in quiescent cells, consistent with the rapid down-regulation of Ki67 expression within 1-2 days after entering the G0 phase (Miller et al., 2018; Sobecki et al., 2016).

Aurora B kinase has been extensively used as a marker for CBs, however its expression in IBs remains uncharacterized. To further differentiate stable IBs from CBs, we labelled midbody-containing bridges using an anti-citron kinase antibody, which marks midbodies in CBs but not mature IBs (Price et al., 2023) and an anti-anillin antibody, previously reported to label IBs in the Drosophila larval brain (Haglund et al., 2011), (**Fig. 2C, S1B**). We found that 81.33% of the bridges were labelled with both Aurora B kinase and anillin (**Fig. 2c_1_** and **2D**), 2.78% were only labelled with Aurora B kinase (**Fig. 2c_2_** and **2D**), and 15.89% were only labelled with anillin (**Fig. 2c_3_** and **2D**) (N = 4 mice, 66 images). The absence of anillin or Aurora B kinase in those two groups may reflect differences in the timing or levels of protein expression, potentially making them undetectable at certain stages of cytokinesis (Halcrow et al., 2022; Renshaw et al., 2014). Importantly, all 3 groups express the midbody protein citron kinase suggesting that all those ICs are CBs (**Fig. 2c1-c3 and 2D**). While we cannot fully exclude the presence of IBs due to the lack of well-characterized markers in somatic cells, our data do not support the presence of IBs in the EGL. Together, these findings indicate that CBs alone cannot account for all the ICs previously identified in the EGL (Cordero Cervantes et al., 2023). Moreover, differences in their distribution and molecular profiles relative to those earlier observations suggest the existence of additional forms of intercellular connectivity. This prompted us to investigate whether other forms of connections could be detected in the developing cerebellum.

### Putative TNT-like connections are present in the developing cerebellum

Having established that the ICs observed in the EGL cannot all be accounted for by CBs or IBs, we next attempted to better characterize these structures (**Fig. S1B)**. We developed an approach to visualize fine structures between sparsely labelled GCs. P5 pups were electroporated to transfer a *pCAG-Dendra2-E2A-GFP-Cep55* plasmid in GCs progenitors (**Fig. S3**), leading to cytoplasmic expression of a fusion protein comprising Dendra2, GFP and the midbody marker CEP55. Pups were sacrificed 2 days after electroporation (Korenkova et al., 2024).

Cerebellar slices were immunostained with anti-GFP to enhance the endogenous cytoplasmic GFP and anti-anillin to visualize CBs (**Fig. 3A-3C**). We used anti-anillin rather than anti-Aurora B kinase, as it identifies a greater number of CBs (**Fig. 2D**). Anti-citron kinase and anti-Cep55 antibodies were not targeted because their punctate staining made specific labelling difficult to distinguish from background. In several cases, we observed pairs of cells that appeared to be linked by a putative bridging structure or a cellular protrusion, compatible with the presence of an IC, although we can not exclude the possibility that the labeled GC processes were merely in close contact with neighboring cell. Some of these potential connections were marked by Anillin, identifying them as CBs. However, among the putative GC connections visualized by GFP staining, 77% did not express anillin (**Fig. 3D**; N = 3 mice, 83 images, 36 unlabeled connections and 11 CBs), suggesting that the majority of them are unlikely to be CBs and therefore may correspond to putative TNT-like structures. This proportion closely matches that reported in gastrula zebrafish embryo, where TNT-like connections, which were also compared to CBs, account for 81% of ICs (Korenkova et al., 2024). However, in contrast to zebrafish, where most TNT-like connections exceeded 5 µm in length, allowing length-based criteria to distinguish CBs/IBs from TNT-like connections, we observed only a modest size difference. Although some putative TNT-like connections extended beyond 5 µm, their mean length was 5.38 µm, compared to 2.92 µm for CBs (**Fig. 3E**), indicating that connection type could not be reliably distinguished by length alone. To directly observe their formation, we performed time-lapse imaging on acute brain slices, capturing a putative TNT-like connection developing via a filopodia-driven mechanism, similar to previously described *in vitro* (Korenkova et al., 2020) (**Fig. 3F-3F’, S3B**). We observed protrusions extending from each cell (white arrowheads), with an apparent connecting bridge between the two cells fully forming within 20 minutes and persisting for at least 10 minutes. This temporal persistence suggests that they are not filopodia, which are much more dynamic structures that protrude and retract within seconds. However, in the absence of functional evidence for cargo exchange, we cannot rule out the possibility that some of the observed ICs correspond to close-ended cell processes or to cytonemes (Korenkova et al., 2020), which are commonly observed during development in *Drosophila* and zebrafish. To assess their open-ended nature and functionality, we attempted to photoconvert the cytoplasmic protein Dendra2 in a single cell and monitor its transfer to the connected cell. However, due to the close apposition of the two cells and light refraction caused by tissue thickness, photoconversion resulted in Dendra2 activation in both cells, precluding definitive interpretation.

### Characteristics of putative TNT-like connections in the EGL from brain slice

To complement the experimental description of those connections and determine whether they occur between clonally or non-clonally related GC, we used a Brainbow-derived strategy. *Nestin-Cre^ERT2+/-^* mice in which tamoxifen induces CRE expression only in neural progenitors were crossed with *Cytbow^+/+^* mice (Loulier et al., 2014) in which mTurquoise, eYFP and tdTomato are stochastically expressed following CRE-mediated recombination. Tamoxifen was administered from P0 to P3 (**Fig. 4A**; see Methods) and resulted in stochastic expression of one of these three fluorescent proteins in GCs, or combinations of two due to the homozygous *Cytbow^+/+^* genotype (**Fig. S4A**). Since these fluorescent reporters are genetically encoded and stably inherited through cell division, they enabled inference of putative clonal relationships among GCs based on color. Although, in principle, neighboring non-clonally related cells could express the same color, sparse labeling minimized this likelihood. Accordingly, cells sharing the same color were considered putatively clonally related, whereas cells labeled with different colors were classified as non-clonally related, as they cannot originate from the same cell division (**Fig. 4A)**. Pups were sacrificed at P7. Similar to the *pCAG-Dendra2-E2A-EGFP-Cep55* plasmid electroporation approach, this strategy enabled sparse labelling of GCs and the identification of putative connections in the densely populated EGL, particularly in lobule X (**Fig. S4B**). In this context, connections between non-clonally related cells (typically of different colors), which cannot result from cell division, may potentially correspond to de novo TNT-like connections. However, given the microscopy limitations, we cannot rule out the possibility that some of these structures in fixed samples are filopodia-like extensions fully spanning two cells. Endogenous fluorescence signals were amplified to optimize optical visualization of these thin structures (**Fig. S4C-E**). The tdTomato signal was enhanced using an anti-tdTomato antibody, while eYFP was amplified using an anti-GFP antibody which also detected mTurquoise2, and to a lesser extent eBFP2 (**Fig. S4C-D**). Importantly, because anti-GFP amplifies both mTurquoise2 and eYFP, these two reporters were not separable within the amplification channel. Nonetheless, given the *Cytbow* design and the sparse labelling, the probability that a mTurquoise2^+^ GC is physically connected to a neighboring eYFP^+^ GC remained limited, minimizing the likelihood of overlooking connections between non-clonally related GCs detected by the anti-GFP antibody. In addition, cerebellar slices were also labelled with an anti-anillin antibody (**Fig. 4A)** to detect CBs between labelled GCs (**Fig. S4E**). We identified both clonally related (87%; **Fig. 4B, 4D, 4H, S4F, S4D**) and non-clonally related (13%; **Fig. 4C, 4D**) GCs connected via putative TNT-like connections (N = 3 mice, 200 images, 27 pairs of putatively clonal-GCs and 4 pairs of non-clonally related GCs). Overall, 8.89% of labelled GCs were connected via putative TNT-like connections (**Fig. 4E**; number of TNT-like connections normalized to the total number of labelled cells).

Among connected cells, 79% possessed a single TNT-like connection (53 out of 67 GCs; **Fig. 4B, F, H, Fig. S4F**) while 21% had multiple connections (14 out of 67 GCs; **Fig. 4C, F, S4d_1_**), a feature previously observed *in vitro* (Sáenz-de-Santa-María et al., 2023). Notably, these connections originated from either soma or neurites, comparable to TNTs described *in vitro* (Tardivel et al., 2016; Vargas et al., 2019). Of the 41 putative TNT-like connections identified, 54% connected GC somas (22/41; **Fig. 4b_1_-b_2_, 4H, S4d_1_, S4F**) 41% connected a soma to a neurite (17/41; **Fig. 4c, 4F purple arrowhead**), and 5% connected two neurites (2/41 connections; **Fig. 4F orange arrowhead**). Additionally, based on the DAPI pattern, we occasionally observed TNT-like connections connecting dividing (identified by compact chromatin) and non-dividing cells (which displayed more diffuse staining and darker nucleolar regions) (**Fig. 4b_1_**), a phenomenon previously reported in developing zebrafish (Korenkova et al., 2024). To further assess the mitotic state of those connected GCs, cerebellar slices were post-stained with anti-PH3 (mitotic marker Phospho-histone H3) to more reliably identify mitotic cells compared to using DAPI staining alone. Additional staining with anti-anillin was used to exclude CBs, and anti-tdTomato was applied to improve the visualization of TNT-like connections (**Fig. 4H**; N = 3 mice, 86 images). We only found putative TNT-like connections between non-dividing cells, suggesting that connections between dividing and non-dividing cells were very rare (**Fig. 4I**). Indeed, although these structures were distributed throughout the EGL, their location varied; some were in the oEGL close to the pia (**Fig. 4b_1_**), others located in the middle of the EGL (**Fig. 4b_2_, S4F**), but the majority were in the iEGL near the ML border, where GCs are mostly postmitotic (**Fig. 4C, 4H, S4d_1_**).

### Conclusion

This study identifies ICs in the EGL of the developing mouse cerebellum, focusing on CBs, IBs, and other types of connections with TNT-like features. We provide evidence for the presence of CBs in the iEGL, whereas IBs do not appear to be present in this region. In addition, we observed diverse morphologies of potential ICs throughout the EGL—thin or thick projections emerging from somata or neurites—which resemble previously reported TNT-like structures *in vitro* and *in vivo* (Chang et al., 2025; Vargas et al., 2019). Notably, our observations suggest that multiple ICs could arise between the same pair of cells. Although GCs predominate in the EGL, our labelling strategy does not exclude the involvement of other cell types in forming these connections (Cadilhac et al., 2021; Palese et al., 2025).

Since all ICs previously identified by ssSEM were open at both ends (Cordero Cervantes et al., 2023), we speculate that some of the connections described here may also be open-ended. However, we cannot exclude the possibility that certain structures represent filopodia-like processes crossing over neighbouring cells or resemble DNTs—closed-end connections—recently reported in the developing mouse cortex (Chang et al., 2025). The lower number of ICs observed here compared to TNT-like structures in zebrafish embryos (35%) (Korenkova et al., 2024) may reflect differences in tissue type, species, developmental timing, or methodology. Fixation and immunostaining procedures are known to disrupt fragile TNTs (Cordero Cervantes & Zurzolo, 2021; Delage et al., 2016; Matejka et al., 2024), and the use of cytoplasmic rather than membrane-targeted reporters may have prevented full visualization of very thin ICs, potentially leading to an underestimation of their frequency.

Overall, our findings are consistent with previous connectomic analyses of the developing mouse cerebellum (Cordero Cervantes et al., 2023), revealing unusual connections between neural cells and excluding a cytokinetic origin for these structures. Our observations are compatible with the presence of TNT-like connections in the EGL that could contribute to cerebellar development by enabling targeted communication before synapse formation (Zurzolo, 2021), coordinating granule-cell migration (Teddy & Kulesa, 2004), or influencing cerebellar organization and synaptogenesis (Kim et al., 2023). Future studies integrating functional analysis with high-resolution live imaging will be essential to determine the nature, dynamics, and developmental roles of these structures in cerebellar circuit assembly.

### Limitations of the Study

Although we identified potential ICs based on 3D morphology, this study could not assess their functional properties. Additional validation is required to determine whether these structures correspond to true TNTs capable of molecular exchange or instead represent cytonemes or filopodia-like protrusions in close proximity to neighboring cells. Functional approaches, including live imaging to track their dynamics and cargo transfer, as well as genetic or pharmacological disruption of their formation, will be critical to establish their role in cerebellar circuit assembly. Our observations, together with our previous ssSEM-based connectomic study (Cordero Cervantes et al., 2023), show that some of these ICs lack cytokinetic and intercellular-bridge markers (Halcrow et al., 2022), arise between non-dividing cells, and appear as thin processes linking distant granule cells, features compatible with TNTs. However, current imaging resolution limits prevent definitive assessment of open-ended continuity, and we cannot exclude the possibility that some structures correspond to transient or incidental protrusions. Functional assays used to study TNT-like structures in other systems, including zebrafish embryos (Korenkova et al., 2024) and interpericyte TNTs (Alarcón-Martínez et al., 2020), have not yet been applied in the developing cerebellum. The lack of specific TNT markers in neural tissue also limits discrimination from other IC types. In addition, the developmental timing of TNT-like connection formation remains unclear; approaches combining pulse–chase labelling with midbody or TNT-associated markers may help define when these structures emerge relative to granule-cell maturation.

Addressing these limitations will be essential to determine whether TNT-mediated communication contributes to cerebellar development and whether its dysregulation may be relevant to neurodevelopmental disorders or cerebellar malformations.

## Acknowledgments

The authors would like to acknowledge the help of the following Institut Pasteur facilities: HistoPathology Core Facility, Photonic Bioimaging platform and the Image Analysis Hub. The authors would like to thank all members of the Zurzolo group for fruitful discussions, with special thanks to Thea Chrysostomou for helping to implement postnatal electroporation at Institut Pasteur and Olga Korenkova for her technical and scientific feedback.

This work was supported by a dedicated grant called Big Brain Theory Program co-funded by Institut Pasteur and Paris Brain Institute to CZ and LC, the Inception program (Investissement d’Avenir grant) ANR-16-CONV-0005, and Equipe FRM - EQU202103012630 ANR AAPG2023 UnProSec 2024-2027, FRM “MND202310017892” 2023-2027 to CZ, Conseil régional d’Ile-de-France, DIM ELICIT and “Financement de Fin de thèse” de la FRM to GK.

## Author contributions

Conceptualization: MR, CZ and LC. Methodology: MR, SL, GK, CZ, LC, FD and JL. Investigation: MR, GK, GV, NDM. Formal analysis: MR, GV. Writing—original draft: MR. Writing—review and editing: MR, GV, SL, CZ, LC, GK, JL, FD. Funding acquisition: CZ, LC

