## Supplemental figure 1-4 for "Deciphering Intercellular Communication in the Cerebellar External Granule Layer: Insights into Non-Classical Connections in Neural Development"

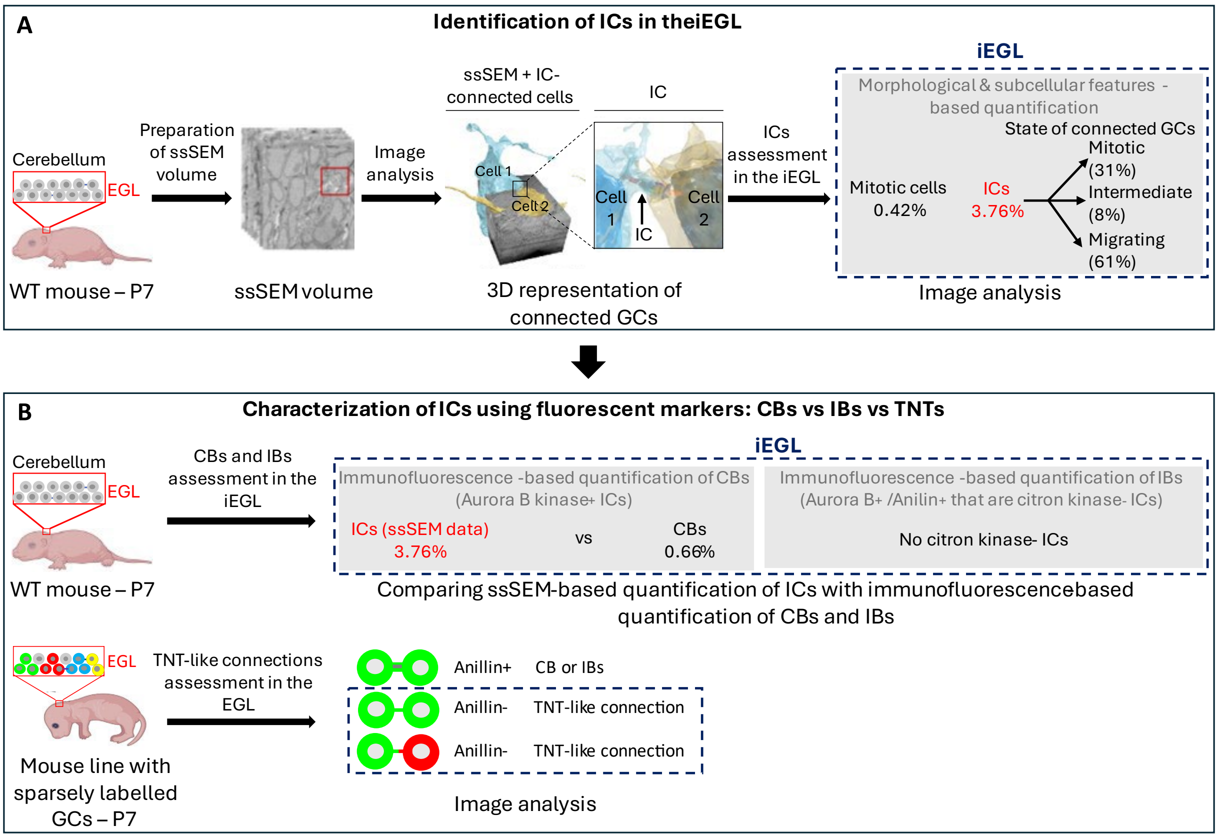


**Fig. S1. Identification of ICs in the developing EGL and assessment of their nature**

(A) Schematic summary of previous work (Cordero et al., 2023) identifying ICs within the iEGL using ssSEM with evidences that some could be TNT-like connections. (B) Schematic overview of the current study, which aims to confirm that a subset of these ICs are neither CBs nor IBs.

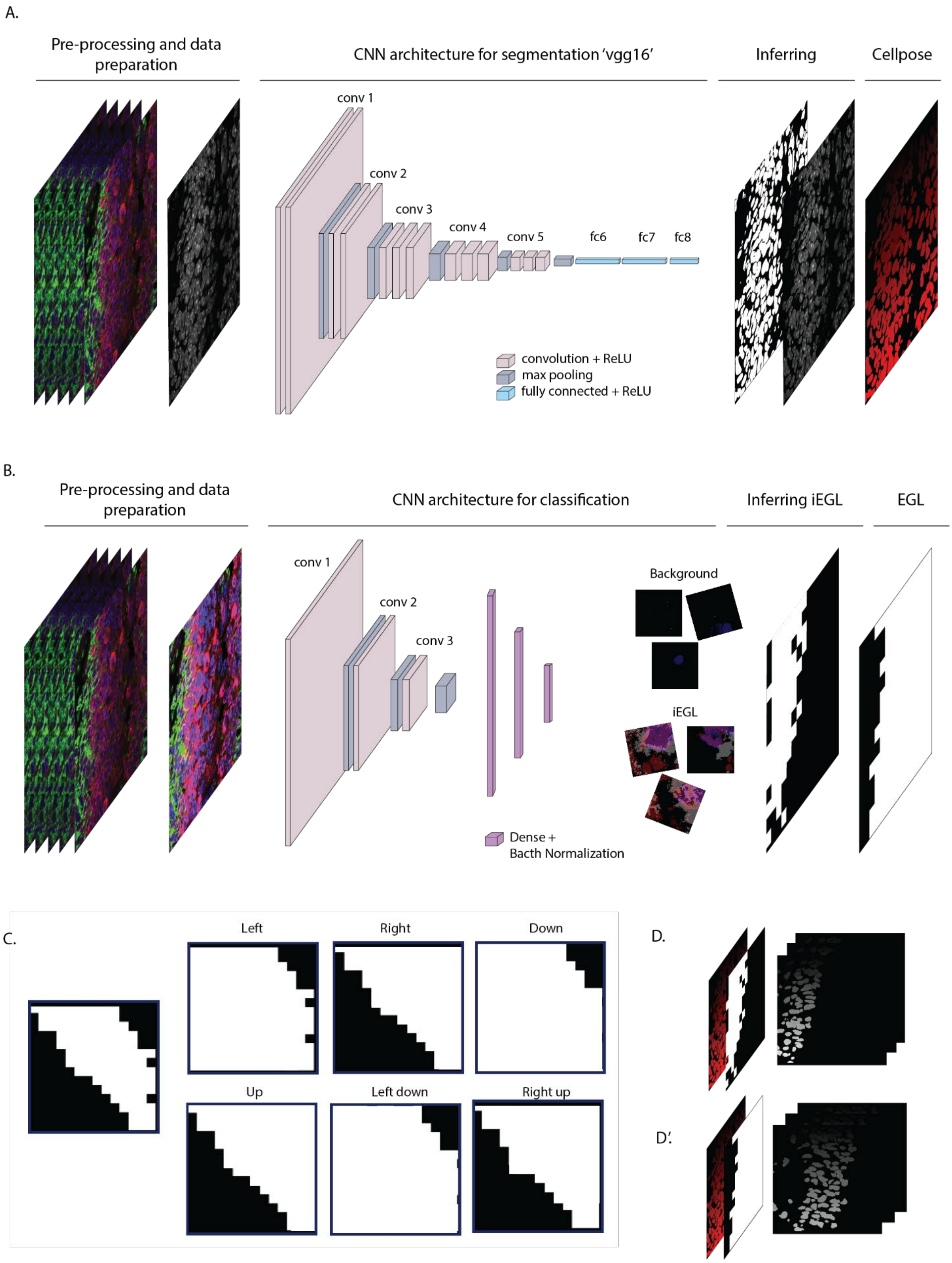
A.
Pre-processing and data preparation
CNN architecture for segmentation 'vgg16'
Inferring
Cellpose
conv 1
conv 2
conv 3
conv 4
conv 5
fc6
fc7
fc8
convolution + ReLU
max pooling
fully connected + ReLU
B.
Pre-processing and data preparation
CNN architecture for classification
Inferring iEGL
EGL
conv 1
conv 2
conv 3
Background
iEGL
Dense +
Bacth Normalization
C.
Left
Right
Down
Up
Left down
Right up
D.
D'.

**Fig. S2. Image processing in the EGL.**

(A) Schematic representation of the steps used for nuclei counting pipeline. The DAPI channel is processed through a CNN-based architecture, followed by nuclei segmentation using the Cellpose software. (B) Schematic showing the delineation of the EGL and iEGL borders for nuclei quantification. DAPI, Ki67 and TAG-1 channels are input to the CNN. The DAPI signal is used to define the EGL while Ki67 and TAG-1 delineate the borders of the iEGL. A classification step distinguishes the iEGL from the rest of the EGL, generating a mask specific to these regions. (C) Illustration of how EGL masks are created based on cerebellum orientation in the image. Orientations include “Left”, “Right”, “Down”, “Up”, ”Left down” or “Right up”. All regions in these orientations are considered part of the iEGL. The images primarily contain the EGL and a small portion of the molecular layer (ML), which contains few nuclei, minimizing quantification errors. (D) Visualization of nuclei identified within the defined masks. Only nuclei located within the mask created - representing either the full EGL or the iEGL - are included in the quantification.

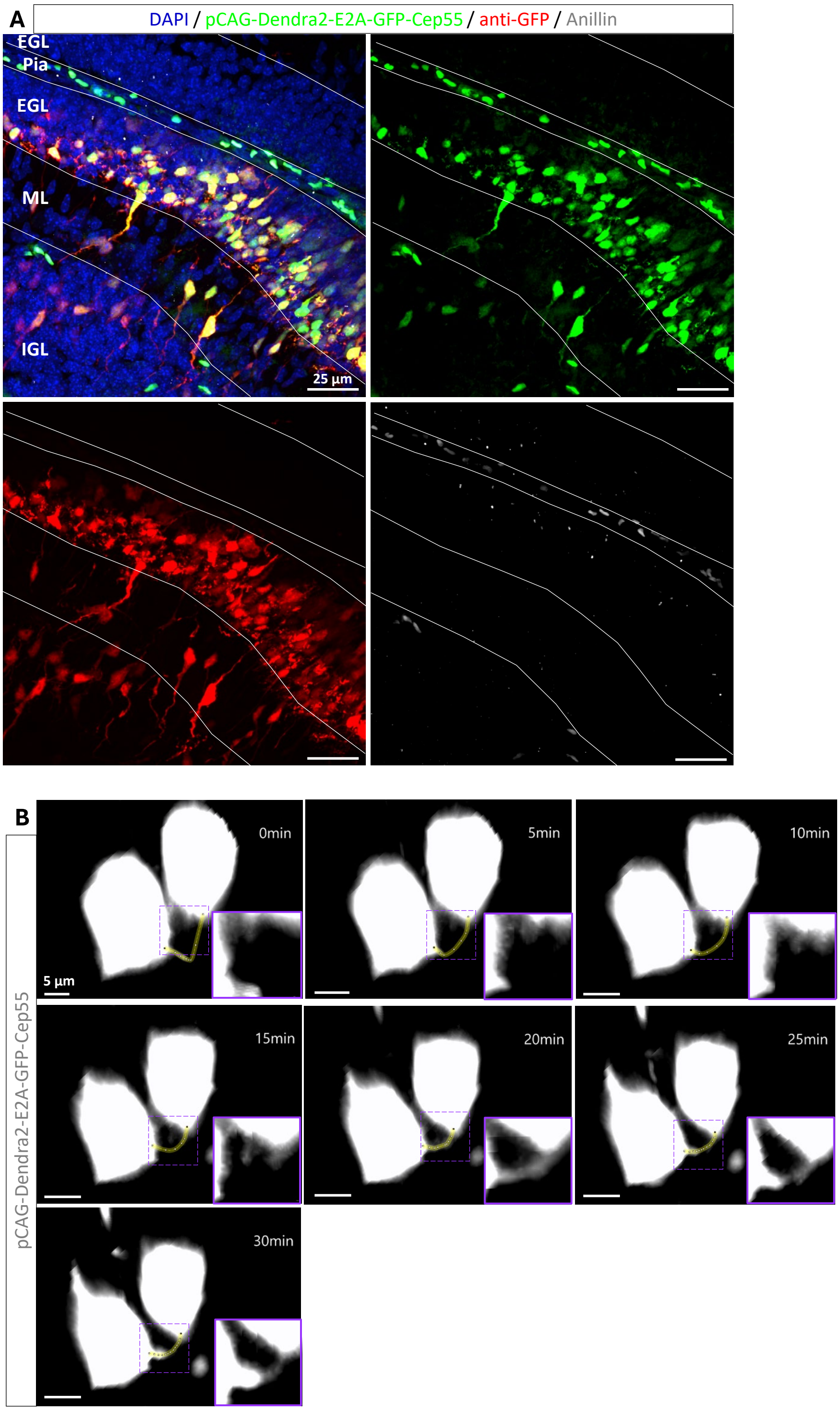


**Fig. S3. Assessment of the expression of the pCAG-Dendra2-E2A-GFP-Cep55 construct**
(A) Representative image of P7 sagittal cerebellar sections electroporated with the pCAG-Dendra2-E2A-GFP-Cep55 construct (green). Sections were stained with anti-GFP (red), anti-anillin (grey) and DAPI, showing that the anti-GFP antibody labels the cytoplasm rather than being exclusively localized to the midbody of electroporated cells. All the images correspond to z-maximum projections and were taken at 10x magnification. Solid lines delineate the different layers of the cerebellum. (B) Location of the line used to generate the fluorescence intensity profile shown in Fig. 3F'.

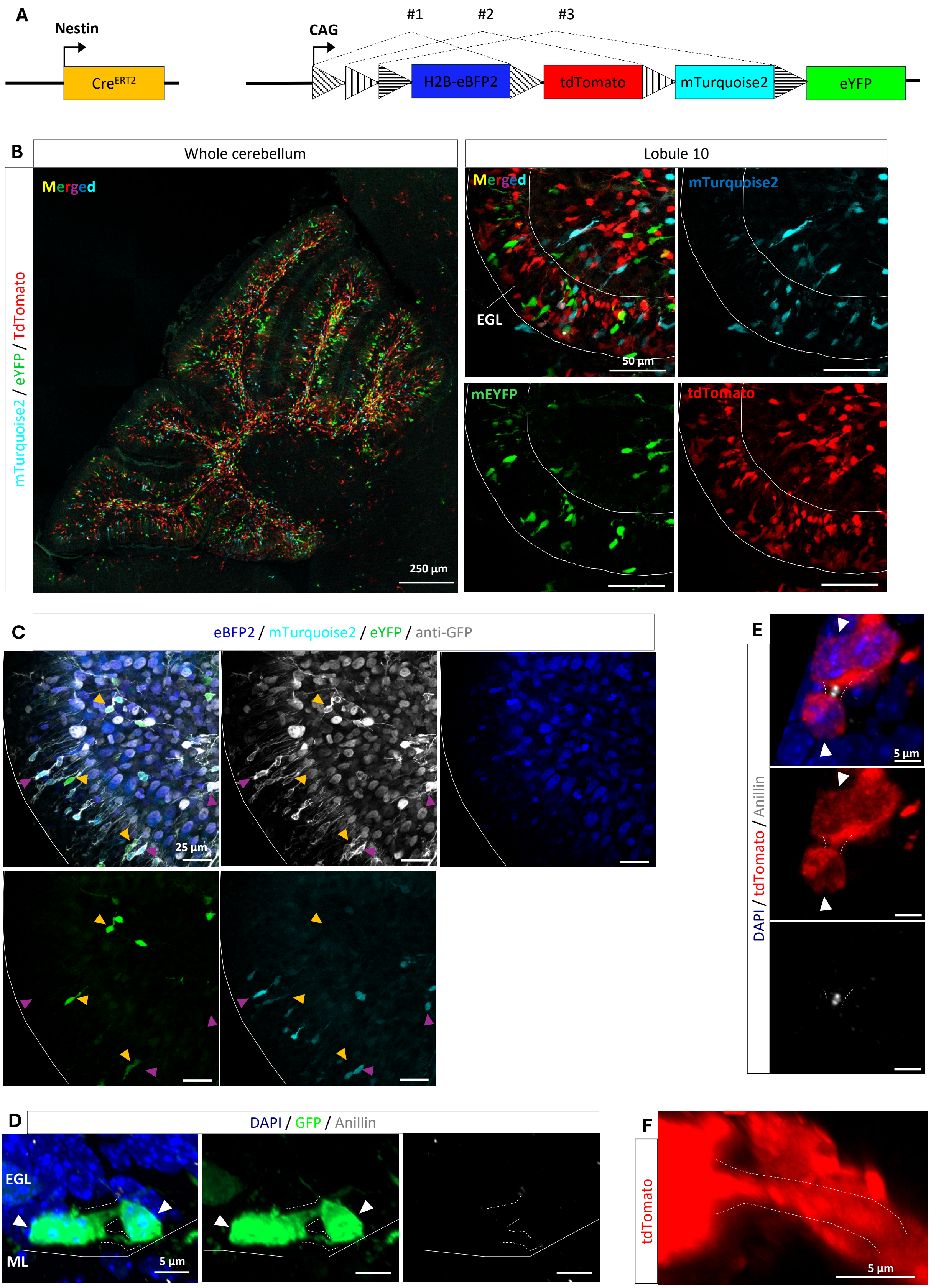
A
Nestin
Cre^ERT2
CAG
#1
#2
#3
H2B-eBFP2
tdTomato
mTurquoise2
eYFP
B
Whole cerebellum
Lobule 10
mTurquoise2 / eYFP / TdTomato
Merged
250 µm
Merged
mTurquoise2
EGL
50 µm
mEYFP
tdTomato
C
eBFP2 / mTurquoise2 / eYFP / anti-GFP
25 µm
E
DAPI / tdTomato / Anillin
5 µm
D
DAPI / GFP / Anillin
EGL
ML
5 µm
F
tdTomato
5 µm

**Fig. S4. Validation of the NestinCre$^{ERT2}$ Cytbow transgenic line**

(A) Schematic representation of the *NestinCre$^{ERT2+/-}$ ; Cytbow$^{+/+}$* transgenic line used to sparsely label cells. This transgenic line expresses nuclear eBFP2 by default. Upon tamoxifen injection, Cre-recombination occurs on the CAG-driven *Cytbow* transgene (Kumamoto et al., 2020). Three mutually exclusive recombination possibilities allow Cre-expressing cells to express a distinct cytoplasmic fluorescent protein: mTurquoise2, mEYFP and tdTomato, in a completely random manner. Since the mice used are homozygous for the *Cytbow* transgene, cells can also express a combination of two reporters. (B) Representative image of P7 sagittal sections showing the endogenous expression of *Cytbow* reporters after tamoxifen injection (P0-P3), in the whole cerebellum, and in the EGL at the level of the lobule 10 where its expression is more pronounced. (C) Labelling with anti-GFP (grey) confirming that it recognizes eBFP2 (blue), mTurquoise2 (Turquoise) and eYFP (green) reporters. Magenta arrowheads highlight mTurquoise2$^{+}$ cells, while orange arrowheads highlight eYFP$^{+}$ cells. (D) Labelling with anti-GFP (green), anti-anillin (grey) and DAPI staining (blue), showing that the anti-GFP staining helps identify TNT-like connections (anillin$^{-}$). The pair of GFP$^{+}$ cells are located at the ML border. The white arrowheads indicate connected cells. Dashed lines delineate TNT-like connections. (E) Labelling of EGL with anti-tdTomato (red), anti-anillin (grey) and DAPI staining (blue). The white arrowheads indicate tdTomato$^{+}$ cells connected via a CB (anillin$^{+}$ connection). Dashed lines delineate the CB structure. (F) Lateral view of connected cells shown in Fig. 2 ($b_2$) showing the TNT-like connection in a different angle. (B-C) images correspond to z-maximum projections. (C-F) images correspond to 3D views. Images were taken at 20x (B) and 63x (C-E) magnification. Solid lines delineate the EGL. Dotted lines delineate the connections.
